# Cocaine receptor identified as BASP1

**DOI:** 10.1101/2020.11.23.392787

**Authors:** Maged M. Harraz, Adarsha P. Malla, Evan R Semenza, Maria Shishikura, Yun Hwang, In Guk Kang, Young Jun Song, Adele M. Snowman, Pedro Cortes, Solomon H. Snyder

## Abstract

Cocaine is a behavioral stimulant with substantial abuse potential related to its positively rewarding actions ^1,2^. Cocaine inhibits the reuptake inactivation of neurotransmitters such as dopamine, serotonin, and norepinephrine at high nanomolar to low micromolar concentrations ^2^. There is evidence for substantially more potent influences of cocaine. For instance, Calligaro and Eldefrawi reported binding of [^3^H]cocaine to brain membranes with a dissociation constant of about 16 nM ^3^. At 10 nM concentration, cocaine elicits environmental place conditioning in planarians ^4^. Furthermore, 1nM cocaine enhances dopamine D2 receptor agonist-mediated signaling ^5^. Inhibition of amine reuptake by cocaine is substantially less potent than some of these high affinity actions. Thus, evidence for a specific, high affinity receptor for cocaine that mediates its behavioral actions has been lacking. We now report high affinity binding of cocaine to the membrane-associated brain acid soluble protein-1 (BASP1) with a Kd of 7 nM. Knocking down BASP1 in the striatum inhibits [^3^H]cocaine binding to striatal synaptosomes. Depletion of BASP1 in the nucleus accumbens diminishes locomotor stimulation, acquisition, and expression of locomotor sensitization to cocaine. Our findings indicate that BASP1 is a pharmacologically relevant receptor for cocaine and a putative therapeutic target for psychostimulant addiction.

## Introduction

Several strategies have been developed to identify targets of small molecule ligands including genetic ^6^, computational ^7^ and biochemical ^8^ approaches. A typical biochemical method relies on affinity purification by immobilizing small molecule ligands as bait on solid supports. Reconstitution of the ligand receptor binding is performed *ex vivo* by running tissue/cell lysates on the small molecule affinity column. After washing unbound material, high concentrations of the small molecule elute bound proteins whose identity is determined by mass spectrometry. We modified this approach to preserve the native environment for ligand receptor binding by binding the ligand to its target(s) in live cells. Following cell lysis, ligand-receptor complexes were isolated by immunoprecipitation.

### Identification of cocaine binding proteins

We pulled down cocaine-binding proteins using an antibody against cocaine. We overexpressed the wild type dopamine transporter (WT-DAT) or L104V-F105C-A109V triple mutant DAT in HEK 293 cells. This mutant was previously reported to be insensitive to cocaine ^9,10^. We incubated cocaine with live cells, then performed the pull-down experiment. Anti-cocaine immunoprecipitation pulls down WT-DAT but not the L104V-F105C-A109V triple mutant DAT only in the presence of cocaine **(Fig. 1a)**. We treated primary cortical neuronal cultures with 100 nM cocaine and incubated the lysates with cocaine antibodies to identify cocaine binding proteins which were eluted with 10 nM cocaine **(Fig. 1b)**. Mass spectrometry analysis reveals six mouse proteins pulled down following treatment with cocaine but not saline. To exclude common contaminants, we checked the frequency of finding these proteins in the Contaminant Repository for Affinity Purification (CRAPome) ^11^ database. Beta actin seems to be a non-specific finding since its maximum number of spectra and the frequency of its occurrence across 411 experiments are very high (**Supplementary Fig. 1a, b)**. We prioritized the brain acid soluble protein-1 (BASP1) as a putative receptor for cocaine, because it was the only membrane-associated protein in our results **(Fig. 1c, d; Supplementary Fig. 1c)** ^12-16^. BASP1 does not seem to be a common contaminant in affinity pull down/mass spectrometry experiments since its relatively low frequency of occurrence is mostly associated with FLAG tag and magnetic beads (**Supplementary Fig. 1d, e)**.

**Figure 1.**
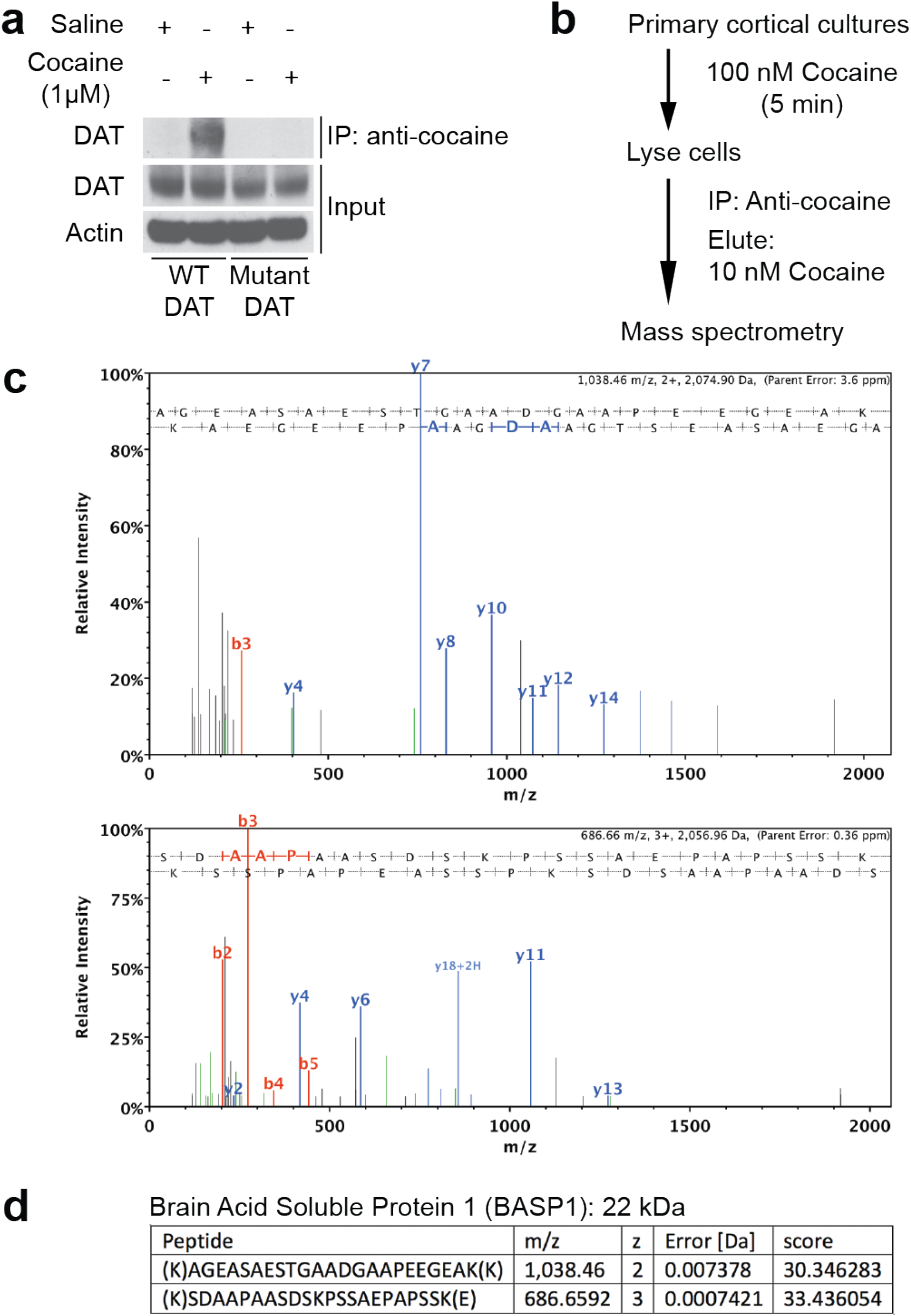
Identification of cocaine binding proteins. **a**, Western blot of DAT overexpressed in HEK 293 cells following immunoprecipitation (IP) with anti-cocaine antibody. Input of DAT and actin from the same sample used in IP serve as loading controls. L104V-F105C-A109V triple mutant DAT is insensitive to cocaine. **b**, A schematic diagram of the protocol we used to identify cocaine binding proteins. **c, d**, MS/MS spectra of the peptides used to identify the brain acid soluble protein-1 (BASP1) in anti-cocaine immunoprecipitates.

### BASP1 is a high affinity cocaine binding protein

We characterized the binding of [^3^H]cocaine to BASP1 protein. [^3^H]cocaine binds overexpressed BASP1 with high affinity, a Kd value of 7 nM **(Fig. 2a; Supplementary Fig. 2a, b)**. [^3^H]cocaine binds to rat striatal synaptosomal fractions with a similar Kd of about 7.9 nM **(Fig. 2b)**. Binding of [^3^H]cocaine to striatal synaptosomes appears to involve BASP1, as depletion of BASP1 by shRNA to about 50% of control levels elicits a 50% decrease in the Bmax of [^3^H]cocaine binding **(Supplementary Fig. 2c, d)**. Depleting BASP1 protein by shRNA treatment diminishes the Bmax with no influence on the Kd of [^3^H]cocaine binding. Similarly, in mouse striatal synaptosomal fractions, depleting BASP1 inhibits [^3^H]cocaine binding **(Fig. 2c, d)**. The decrease of BASP1 protein by shRNA treatment corresponds closely to the reduction of [^3^H]cocaine binding consistent with endogenous BASP1 being responsible for all or the great majority of high affinity [^3^H]cocaine binding. Receptor specificity discriminates substances with varying resemblance to its ligand. We compared diverse cocaine derivatives for their ability to elute [^3^H]cocaine from BASP1 containing membranes. Out of twelve tested compounds, only benztropine and 3β-(p-Fluorobenzoyloxy)tropane (3-p-FBT) elute [^3^H]cocaine from BASP1 containing membranes. Other closely related compounds such as the major cocaine metabolite benzoylecgonine does not dissociate [^3^H]cocaine from BASP1 **(Fig. 2e; Supplementary Fig. 2e)**. These findings indicate that BASP1 is a selective, high affinity cocaine binding protein.

**Figure 2.**
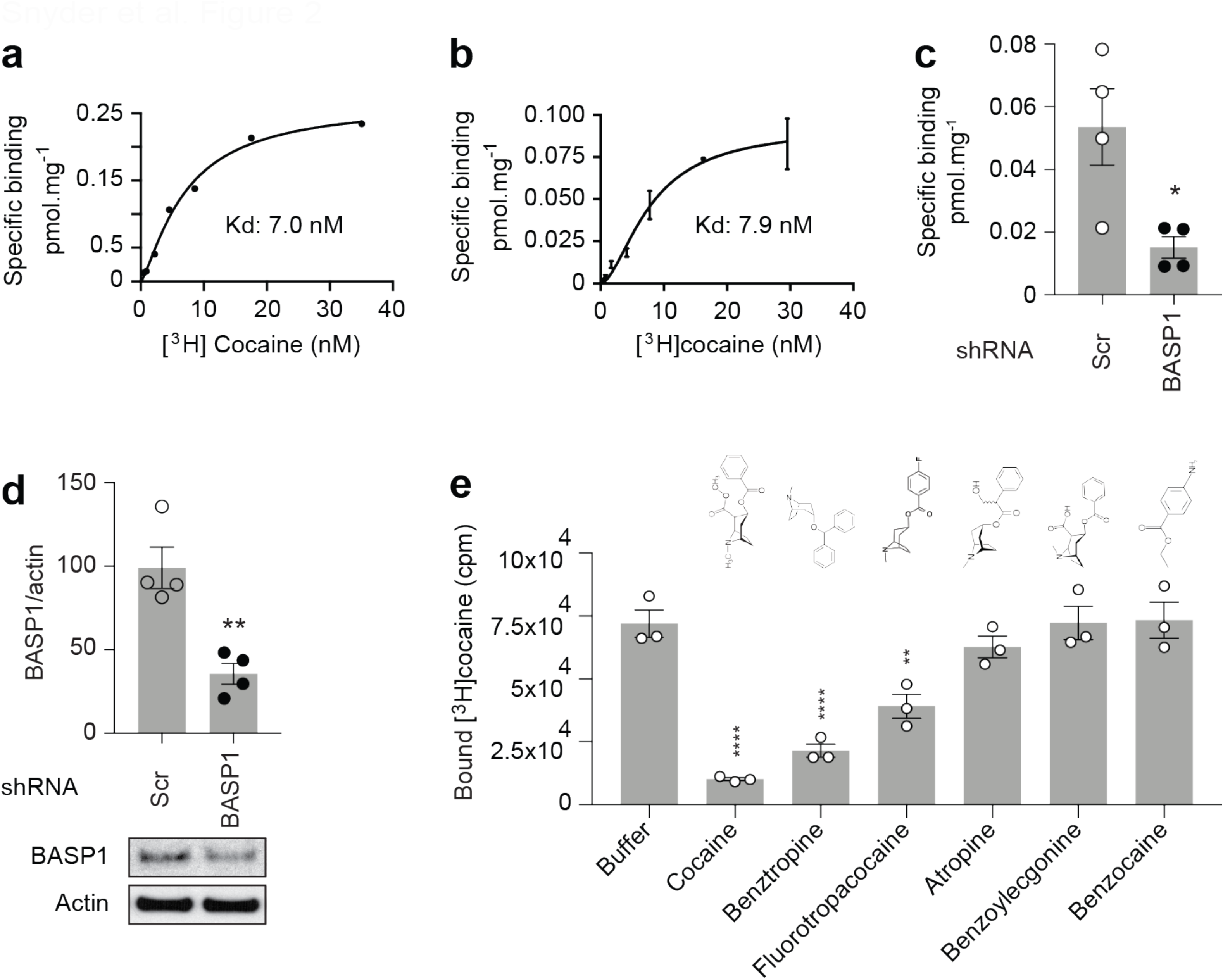
BASP1 is a high affinity cocaine binding protein. **a**, Saturation binding curve for [^3^H]cocaine and BASP1 overexpressed in HEK-293 cells and anchored to the membrane with a PDGF transmembrane domain. Membrane fraction is used for the saturation binding assay. Kd = 7.0 nM, R square = 0.99, specific binding with Hill slope. **b**, Saturation binding curve for [^3^H]cocaine to rat striatal synaptosome fraction. Kd = 7.9 nM, R square = 0.97, *n* = 2, Error bars = +/-SD, specific binding with Hill slope. **c**, AAV2 delivered shRNA to BASP1 reduces specific binding of [^3^H]cocaine to mouse striatal synaptosome fraction. *n* = 4, * (*P* = 0.023), two-tailed *t* test. Error bars = +/-SEM. **d**, Western blot of BASP1 in NAc preinjected with AAV2 that encodes scrambled (Scr) or BASP1 shRNA. The westerns are for the same samples used in panel **c**. Quantification of western blot band intensity of BASP1 normalized to actin is shown. *n* = 4, ** (*P* = 0.0038), two-tailed *t* test. Error bars = +/-SEM. **e**, Specific binding of [^3^H]cocaine to membrane fraction of HEK 293 cells overexpressing BASP1 following elution with indicated structural analogues/derivatives of cocaine. *n* = 3, one-way ANOVA, *P* < 0.0001, Bonferroni’s multiple comparisons test, **** (*P* <0.0001), ** (*P* = 0.0056). Error bars = +/-SEM.

### BASP1 mediates the locomotor stimulation by cocaine

We explored the influence of BASP1 depletion in the nucleus accumbens (NAc) of intact mice utilizing shRNA incorporated into an adeno-associated viral vector (AAV2) **(Fig. 3a)**. This shRNA treatment elicits a 50% depletion of BASP1 in the nucleus accumbens **(Fig. 3b)**. We evaluated the effect of BASP1 depletion upon the behavioral effects of cocaine monitored as enhancement of the horizontal locomotor activity by a single injection of cocaine (20 mg/kg) **(Fig. 3c)**. BASP1 depletion without cocaine treatment does not affect spontaneous locomotor activity of mice **(Fig. 3c, d)**. In control animals receiving scrambled shRNA cocaine substantially enhances locomotor activity. BASP1 shRNA treatment reduces the stimulatory actions of cocaine by about 50%, which corresponds to 50% reduction of endogenous BASP1 protein **(Fig. 3c, d)**. BASP1 depletion inhibits the cocaine-induced peripheral **(Supplementary Fig. 3a)** but not central locomotor activity **(Supplementary Fig. 3b)**. During the peak cocaine-induced locomotor activity, BASP1 depletion prevents the reduction by cocaine of vertical/rearing activity **(Supplementary Fig. 3c)**.

**Figure 3.**
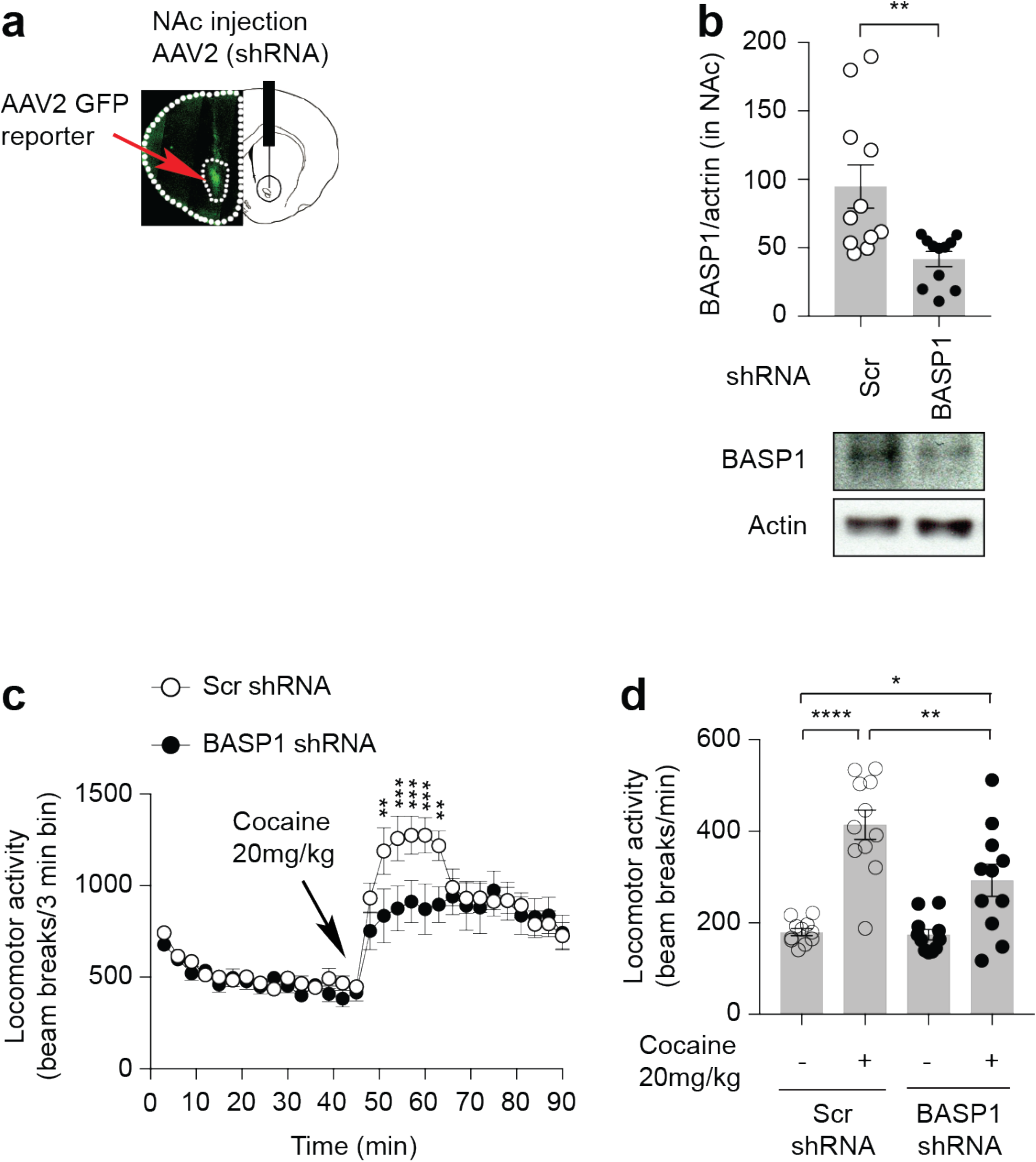
BASP1 mediates the locomotor stimulant effect of cocaine. **a**, GFP reporter confirmation of NAc targeting by stereotaxic injection. **b**, Western blot of BASP1 in NAc preinjected with AAV2 that encodes scrambled (Scr) or BASP1 shRNA. The westerns are from the same samples used in the behavioral assay (**c, d**). Quantification of western blot band intensity of BASP1 normalized to actin is shown. Samples with more than 66.6% of the average control levels of BASP1 (as determined by western blot) were considered unsuccessful knockdown and excluded from the open field test data analysis. *n* = 11 per group, ** (*P* = 0.0048), two-tailed *t* test. Error bars = +/-SEM. **c**, Total locomotor activity of mice in the open field test treated with cocaine preinjected with AAV2 scrambled (Scr) or BASP1 shRNA. *n* = 11 mice/group, two-way ANOVA, ** (*P* = 0.0013 and 0.0034 respectively), *** (*P* = 0.0005, 0.0010, and 0.0003 respectively). Error bars = +/-SEM. **d**, Peak locomotor activity of mice in the open field test. *n* = 11 mice/group, one-way ANOVA, *P* < 0.0001, Bonferroni’s multiple comparisons test, **** Scr shRNA group saline vs cocaine treatment (*P* < 0.0001), ** Scr shRNA + cocaine vs BASP1 shRNA + cocaine (*P* = 0.0078), * Scr shRNA saline vs BASP1 shRNA + cocaine (*P* = 0.015). Error bars = +/-SEM.

### BASP1 regulates the acquisition and expression of locomotor sensitization to cocaine

Repeated administration of cocaine results in a progressive and long-lasting increase in the locomotor stimulant effect triggered by a later drug injection/challenge. This phenomenon, known as locomotor sensitization to cocaine, likely underlies some aspects of cocaine abuse ^17^. We utilized the same *in vivo* knockdown approach to test the effect of BASP1 depletion in NAc on the locomotor sensitization to cocaine in mice. In control mice injected with scrambled shRNA, repeated administration of cocaine (15 mg/kg) elicits locomotor sensitization **(Fig. 4a, b; Supplementary Fig. 4a-g)**, which is still expressed a week following the last consecutive dose of cocaine (day 14) **(Fig. 4a-c)**. BASP1 depletion in NAc markedly reduces the acquisition of locomotor sensitization in mice following repeated injections of cocaine **(Fig. 4a, b; Supplementary Fig. 4a-g)**. Furthermore, depletion of BASP1 in NAc abolishes the expression of locomotor sensitization to cocaine on day 14 **(Fig. 4b, c)**. These findings suggest that BASP1 mediates both acute and chronic behavioral stimulant actions of cocaine.

**Figure 4.**
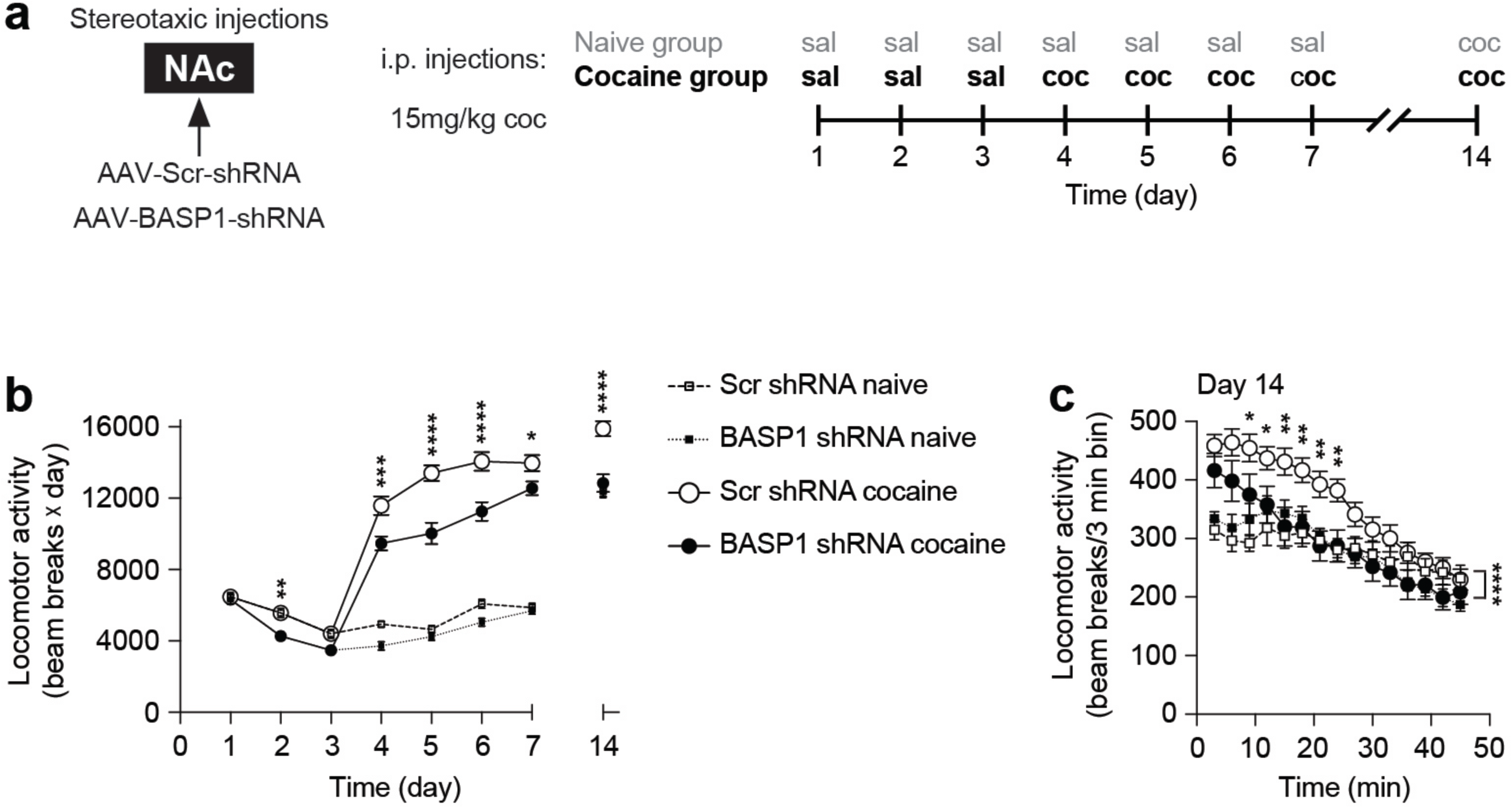
BASP1 regulates the acquisition and expression of locomotor sensitization to cocaine. **a**, A schematic diagram of the locomotor sensitization experimental protocol. **b**, Total open field locomotion during 45 min each day after cocaine injection (15 mg/kg). Days 1-3: baseline, days 4-7: acquisition of locomotor sensitization, day 14: cocaine challenge (15 mg/kg). Scr shRNA naive group: *n* = 7, BASP1 shRNA naive group: *n* = 9, Scr shRNA cocaine group: *n* = 16, BASP1 shRNA cocaine group: *n* = 13. Two-way ANOVA, *P* < 0.0001, Bonferroni’s multiple comparisons test, * (*P* = 0.03), ** (*P* = 0.008), *** (*P* = 0.0002), **** (*P* < 0.0001). Error bars = +/-SEM. **c**, Cocaine challenge day open field locomotor activity. All groups are injected with cocaine (15 mg/kg) right before the test. Scr shRNA naive group: *n* = 7, BASP1 shRNA naive group: *n* = 9, Scr shRNA cocaine group: *n* = 16, BASP1 shRNA cocaine group: *n* = 13. Two-way ANOVA, Bonferroni’s multiple comparisons test, **** (*P* < 0.0001) Scr shRNA cocaine vs BASP1 shRNA cocaine, * (*P* = 0.047 and 0.048 respectively), ** (*P* = 0.0013, 0.0067, 0.0027, and 0.0098 respectively). Error bars = +/-SEM.

## Discussion

Our findings indicate that BASP1 (also known as NAP□22 and CAP□23) is a pharmacologically relevant cocaine receptor. Thus, BASP1 is the principal membrane associated protein interacting with cocaine in rodent brain and binds cocaine with notably high affinity (Kd, 7 nM). Depleting BASP1 leads to a corresponding decrease of [^3^H]cocaine binding indicating that BASP1 accounts for all or the great majority of high affinity [^3^H]cocaine binding interactions in the brain. The enhancement of locomotor activity by cocaine is reduced by depleting NAc BASP1 with shRNA. Furthermore, BASP1 depletion in NAc reduces the acquisition and expression of locomotor sensitization to cocaine. Earlier Calligaro and Eldefrawi^3^ reported high affinity binding of [^3^H]cocaine in rat brain membranes, whose properties closely resemble BASP1.

BASP1 is a membrane associated protein whose myristoylation links it to brain membranes. It is enriched in axon terminals and regulates neurite outgrowth, maturation of the actin cytoskeleton, and organization of the plasma membrane ^12,13,18,19^. However, its exact biochemical and behavioral functions have not heretofore been elucidated. BASP1 is phosphorylated by protein kinase C and physiologically binds calmodulin ^20,21^. It is associated with cholesterol, a major component of lipid rafts in the brain ^22^. Some evidence suggests that BASP1 regulates the formation of lipid rafts ^23,24^. Despite its small size, BASP1 induces cation□selective currents across negatively□charged planar lipid bilayers ^25^. BASP1 is functionally related to the growth-associated protein-43 (GAP-43) or neuromodulin and the myristoylated alanine-rich C-kinase substrate (MARKS) which are enriched in presynaptic terminals. Specific interactions of GAP-43, MARKS and BASP1 with phospholipids and actin contribute to the regulation of neurotransmitter release, axonal growth cone guidance, and synaptic plasticity ^26^.

BASP1 also has actions outside of the brain. For instance, in kidney derived cells BASP1 binds to Wilms tumor factor for which it serves as a transcriptional co-repressor ^15,16^. Furthermore, BASP1 seems to act as a tumor suppressor by inhibiting myc-induced oncogenesis ^27^.

In summary, BASP1 displays properties consistent with its serving as a cocaine receptor. Accordingly, drugs that act selectively upon BASP1 might facilitate or antagonize actions of cocaine. BASP1 may physiologically regulate mood and alertness.

## Materials and Methods

### Reagents

Neurobasal-A medium (no glucose, no sodium pyruvate), B-27 supplement and B-27 supplement minus antioxidants were purchased from Life Technologies (Grand Island, NY). Cocaine hydrochloride, atropine, benzoic acid, benztropine, ecgonine, lidocaine, scopolamine and glutathione agarose were purchased from Sigma-Aldrich (St. Louis, MO). Anisodamine, benzocaine, benzoylecgonine, benztropine, 3-p-FBT (hydrochloride), procaine, and tropine were purchased from Cayman Chemical (Ann Arbor, Michigan). Protein A/G plus-agarose was purchased from Santa Cruz Biotechnology (Paso Robles, CA). [^3^H]cocaine was purchased from PerkinElmer Life Sciences (Boston, MA). FLAG-DAT plasmid (pcihygroflagsynhDAT) was a gift from Jonathan Javitch, Columbia University (Addgene plasmid # 19990) ^28^. PDGF-TMD-GFP plasmid (pFU-no toxin-PE) was a gift from Ines Ibanez-Tallon (Addgene plasmid # 24149) ^29^.

### Cloning

A FLAG tag followed by the PDGF transmembrane domain (TMD) were cloned to the 5’ end of BASP1 removing its ATG start codon to create a FLAG-PDGF-TMD-BASP1 fusion protein. BglII-AsiSI restriction sites were used to insert the FLAG-PDGF-TMD fragment. The FLAG-PDGF-TMD fragment was derived from pFU-no toxin-PE Addgene Plasmid #24149. The correct clone was identified by DNA sequencing to exclude any point mutants followed by overexpression in HEK293 cells, isolation of the membrane fraction and western blot for the FLAG tag.

### Antibodies

Beta Actin-HRP antibody; GeneScript (Piscataway, NJ), catalog number: A00730, clone 2D1D10, lot number: 13B000572. Sheep anti-cocaine antibody was purchased from US Biological Life Sciences (Salem, MA), catalog number: C7505, lot number: L0031710A. Anti-BASP1 antibody used in figure 3 was purchased from Santa Cruz Biotech (Paso Robles, CA), catalog number: sc-66994, clone H-100, lot number: B1715. BASP1 Antibody-C-terminal region (used in figure 2), Aviva Systems Biology (San Diego, CA), catalog number: ARP59932_P050, lot number: QC69877-43224. DAT antibody used for western blots; Santa Cruz Biotech, catalog number: sc-32258, clone 6-5G10, lot number: D0111. ANTI-FLAG antibody; Sigma Aldrich, catalog number: F3165, clone M2, lot number: SLBQ7119V.

### Ligand binding assay

FLAG-tagged PDGF transmembrane domain fused to GFP on its N-terminus (FLAG-TMD-GFP) or to BASP1 on its N-terminus (FLAG-TMD-BASP1), or myc-FLAG fused to BASP1 on its C-terminus (BASP1-myc-FLAG) were overexpressed in HEK293 cells; enrichment at the membrane fraction was verified using subcellular fractionation and western blot. Membrane fractions or striatal synaptosomal fractions were incubated with [^3^H]cocaine for 30 min at room temperature. Then the mixtures were passed through a polyethylenimine (PEI) coated Whatman filter. The radioactivity retained on filters after 4 x washing was monitored using a scintillation counter; Beckman Coulter LS6500 Liquid Scintillation Counter, LS6500 software, Beckman Coulter (Fullerton, CA). Specific binding (total binding - binding in the presence of excess unlabeled cocaine) was analyzed by curve fitting using non-linear regression analysis in order to calculate the Kd and Bmax of cocaine-BASP1 binding.

### HEK-293 cells transfection

HEK-293 cells were transfected using calcium phosphate method. Briefly, plasmid DNA was diluted in 2X HBS buffer containing 1.5 mM disodium phosphate, 12 mM dextrose, 280 mM NaCl, 10 mM KCl, and 50 mM HEPES pH 7.05. Then, 0.25 M calcium chloride was added dropwise to the mix while vortexing on low speed. Transfection mixers were incubated at room temperature for 30 min then added to cells in a drop wise fashion. The media was changed 24 h later. The cells were used 48-72 h after transfection.

### Isolation of membrane fractionation

HEK-293 cells were homogenized in buffer containing 0.25 M sucrose, 10 mM HEPES (pH 7.3), 1 mM EDTA, 0.5 μg/ml antipain, 1 μg/ml leupeptin, 1 μg/ml aprotinin, 1 μg/ml chymostatin and 1 μg/ml pepstatin A. The homogenates were centrifuged at 1,000 x g for 10 min at 4 °C. The supernatants were centrifuged at 3,000 x g for 10 min at 4 °C. The supernatants from the last step were centrifuged at 10,000 x g for 10 min at 4 °C. Resultant supernatants were then loaded into ultracentrifuge SW-55Ti rotor tube and spun at 100,000 x g for 1hr at 4 °C. Resulting pellets (membrane fractions) were resuspended in 10 mM sodium phosphate buffer (with 0.5 μg/ml antipain, 1 μg/ml leupeptin, 1 μg/ml aprotinin, 1 μg/ml chymostatin and 1 μg/ml pepstatin A) to a concentration of approximately 0.5 mg/ml protein.

### Animals

All experiments involving animals were conducted in accordance with the Johns Hopkins Medical Institutions Animal Care and Use Committee guidelines. C57Bl/6j adult mice were purchased from The Jackson Laboratory (Bar Harbor, ME). CD-1 timed pregnant mice and Sprague Dawley male rats were purchased from Charles River Laboratories. Animal handling and procedures were conducted in accordance with the National Institutes of Health guidelines for use of experimental animals and the Johns Hopkins animal care and use guidelines. For sample size determination, we used the Sample Size Calculator at ClinCalc.com.

### Cocaine immunoprecipitation (IP)

Cortical cultures were treated with 100 nM cocaine for 5 min, then washed in ice-cold PBS and harvested on ice in IP buffer containing: 10 mM sodium phosphate pH 7.2, 1% NP-40, 100 mM sodium chloride, 2 mM EDTA, 50 mM sodium fluoride, 200 µM sodium orthovanadate, 0.5 μg/ml antipain, 1 μg/ml leupeptin, 1 μg/ml aprotinin, 1 μg/ml chymostatin and 1 μg/ml pepstatin A. Cell lysates were passed through a 27 gauge syringe ten times, incubated on ice for 15 min then cleared by centrifugation at 3,000 x g for 10 min at 4°C. Equal amounts of total protein were incubated with the sheep anti-cocaine antibody for 45 min at 4°C with rotation. Then, protein A/G plus agarose beads were added to the lysate-antibody mixture and rotated at 4°C for 1 h. The beads were washed 5 times by a series of centrifugation (at 735 x g) and resuspension using TBS buffer. Elution was performed using 10 nM cocaine. The eluates were then mixed with reducing SDS-loading buffer, boiled and separated by SDS-PAGE or sent for mass spectrometric identification.

### Primary cortical neurons (PCNs) culture

Unless otherwise specified, PCNs were isolated from E16-E18 pregnant CD-1 mice. The pregnant mouse was sacrificed by decapitation, then the uterus was dissected out immediately. Working in sterile conditions, the uterus was opened, the pups decapitated, and their brains dissected and placed in dissection media containing DMEM/F12 1:1 supplemented with 10% horse serum. The cerebral cortices were detached from the rest of the brain, and the meninges removed. The cortices were incubated in 0.025% trypsin for 15 min at 37°C. The trypsin was washed with dissection media. The cortices were disrupted into single cell suspension by pipetting up and down 10 times, then strained through a 40 µm sterile mesh. The single cell suspension was cultured in dissection media overnight. Then the media was replaced by PCNs plating media containing neurobasal-A medium (no glucose, no sodium pyruvate) supplemented with 12.5 mM glucose, 2 mM l-glutamine and 2% B-27. On day-4 in vitro (DIV4) the media was changed to PCNs maintenance media containing neurobasal-A medium (no glucose, no sodium pyruvate) + 12.5 mM glucose, 2 mM l-glutamine and 2% B-27 minus antioxidants. Every 3 days thereafter, 50% of the media was replaced with PCNs maintenance media.

### Brain tissue processing

Mice were anesthetized with intraperitoneal injection of sodium pentobarbital (80 mg/kg). The thoracic cavity was exposed, and trans-cardiac perfusion was performed by inserting a needle to the left ventricle and puncturing the right atrium. Phosphate buffered saline (PBS) was used to start the perfusion for 5 min at a rate of 5 ml/min followed by ice cold 4% paraformaldehyde (PFA) in PBS for 30 min (using 3 ml/min rate). The brains were dissected and post-fixed in 4% PFA for 24 h at 4°C then cryo-protected with 30% sucrose in PBS for 24 h at 4°C. Brains were sectioned on a freezing stage sliding microtome into a series of 40 μm sections. ZEISS LSM 800 confocal microscopy was used to image sections. ZEN imaging software (2.3 blue edition service pack 1), Carl Zeiss Microscopy, (Jena, Germany) was used for data collection.

### Stereotaxic injection

Stereotaxic injections were performed as previously described with modifications ^30^. C57 BL6/j mice (age range:10-14 weeks) were anesthetized with 125 mg/kg ketamine, 15 mg/kg xylazine, and 1.5 mg/kg acepromazine. Stoelting’s stereotaxic instrument for mice was used to perform the procedure. The skull was exposed, and burr holes made at the stereotaxic co-ordinates specified below. The nucleus accumbens (NAc) was injected bilaterally using the following stereotaxic coordinates from bregma: + 1.4 mm (anterior/posterior), +/-1.3 mm (medial/lateral), and −4.3 mm (dorsal/ventral). One μl of AAV2 was delivered on each side at a rate of 1 μl/min, followed by 5 min pause at site of injection, 2 min pause at −3.3 (dorsal/ventral), and 2 min pause at −2.3 (dorsal/ventral). For the striatal injections, from bregma: -0.5 mm (anterior/posterior); +/-2.0 mm (medial/lateral); 4 x 1 μl AAV2 injections were delivered on each side at a rate of 1 μl/min followed by a 2 min pause at the following depths (dorsal/ventral) -4.5 mm (1 μl), -4.0 mm (1 μl), -3.5 mm (1 μl), and -3.0 mm (1 μl). Then, 5 min long pauses were made at the following depths (dorsal/ventral) -2.5 mm and -2.0 mm. Confirmation of target site was verified by visualizing GFP expression by confocal microscopy. AAV2 encoding shRNA against BASP1 was purchased from Vector Biolabs (Malvern, PA), AAV2 encoding scrambled (Scr) shRNA was purchased from Vector Biolabs (Malvern, PA) and the University of Iowa Viral Vector Core (Iowa City, IA). Knockdown of BASP1 was verified by western blot. Western blot band intensities were quantified using NIH-image J 2.0.0, NIH, USA.

### Mass Spectral analysis: Proteolysis

After adjusting pH to 8.0 with 4 μl triethylammonium bicarbonate (TEAB) buffer, samples were reduced with 2 μl x 7.5 mg/mL dithiothreitol (DTT) at 60°C for 1 h, alkylated with 2 μl x 18.5mg/mL iodoacetamide in the dark at RT for 15 min. Proteins were proteolyzed with 650 ng trypsin (lyophilized, Promega, www.promega.com) at 37°C overnight. Tryptic peptides were desalted on Oasis u-HLB plates (Waters), then eluted with 65% acetonitrile/0.1%TFA.

### LC/MS/MS analysis

Desalted tryptic peptides were analyzed by liquid chromatography/tandem mass spectrometry (LCMS/MS) on nano-LC-QExative HF in FTFT (Thermo Fisher Scientific, www.thermofisher.com) interfaced with nano-Acquity LC system from Waters, using reverse-phase chromatography (2%-90% acetonitrile/0.1% FA gradient over 75 min at 300 nl/min) on 75 µm x 150 mm ProntoSIL-120-5-C18 H column 5 µm, 120Å (BISCHOFF) http://www.bischoff-chrom.com/hplc-prontosil-c18-h-c18-phasen.html Eluting peptides were sprayed into an QExactive HF mass spectrometer through an 1 µm emitter tip (New Objective, www.newobjective.com) at 2.0 kV. Survey scans (Full MSs) were acquired on Orbi-trap within 350-1800Da m/z using data dependent Top 15 method with dynamic exclusion of 16 s. Precursor ions were individually isolated with 1.6 Da, fragmented (MS/MS) using HCD activation collision energy 27 or 28. Precursor and the fragment ions were analyzed at resolution 120,000 AGC target 3×10^6^, max IT 100 ms and 60000, AGC target 1×10^5^, mx IT200ms, respectively.

### Data Analysis

Tandem MS2 spectra were processed by Proteome Discoverer (v1.4 ThermoFisher Scientific) in three ways, using 3Nodes: common, Xtract (spectra are extracted, charge state deconvoluted, and deisotoped using Xtract option, at resolution 95 K at 400 Da), MS2 Processor. MS/MS spectra from 3Nodes were analyzed with Mascot v.2.5.1 Matrix Science, London, UK (www.matrixscience.com) specifying trypsin as enzyme, missed cleavage 2, precursor mass tolerance 8 ppm, fragment mass tolerance 0.015Da and Oxidation(M), carbamidomethyl c, Deamidation NQ as variable modifications.

### Open field test

Open field activity monitoring was performed in an open field Plexiglas chamber with photocell emitters and receptors to detect locomotor activity using a grid of invisible infrared beams; Photobeam Activity System, SDI PAS-Open Field software, San Diego Instruments, (San Diego, CA). The chambers were enclosed in illuminated opaque boxes and connected to a computer for beam break data collection. Age matched male mice preinjected in NAc with AAV2 encoding Scr or BASP1 shRNA (2 weeks prior to experiment) were placed in the chambers for 45 min to detect baseline activity. After baseline monitoring, mice were given intraperitoneal injections of cocaine (20 mg/kg) or saline and returned to the chambers for 45 min to monitor their locomotor behavior. Cocaine’s effect on peak locomotor activity was calculated based on the 15 min following cocaine injection.

### Locomotor sensitization experiments

Age matched male and female mice were injected in NAc with AAV2 encoding Scr or BASP1 shRNA then randomly divided into naive and cocaine groups. Locomotor sensitization experiments were performed 2 weeks after the stereotaxic injection. The naive group received i.p. saline injection from days 1-7 and a 15 mg/kg cocaine injection on challenge day (day 14). For the cocaine group, mice were injected as follows: d 1-3 i.p. saline; d 4-7 and d 14 i.p. cocaine (15 mg/kg). Mice were immediately tested for locomotor activity for 45 min after each injection in an open field Plexiglas chamber; Photobeam Activity System, SDI PAS-Open Field software, San Diego Instruments, (San Diego, CA).Total beam breaks over 45 min per day were measured.

### Synaptosome fraction preparation

Crude synaptosome fractions were prepared as previously described ^31^ with minor modifications. Briefly, striata derived from male mice or rats were dissected and homogenized in a buffer containing 0.32 M sucrose, 20 mM HEPES (pH 7.4), 0.5 μg/ml antipain, 1 μg/ml leupeptin, 1 μg/ml aprotinin, 1 μg/ml chymostatin and 1 μg/ml pepstatin A. The tissue homogenates were centrifuged at 3000xg for 10 min at 4 °C. The supernatants were centrifuged at 10,000xg for 15 min at 4 °C. The pellets (the crude synaptosome fraction) were resuspended in RIPA buffer (Cell Signaling Technology, Danvers, MA) containing 20 mM Tris-HCl (pH 7.5), 150 mM NaCl, 1 mM Na2EDTA 1 mM EGTA, 1% NP-40, 1% sodium deoxycholate, 2.5 mM sodium pyrophosphate 1 mM b-glycerophosphate, 1 mM Na_3_VO_4_, and 1 μg/ml leupeptin. The crude synaptosome fraction lysates were used for western blot analysis. Western blot was performed following standard procedures except for the dopamine transporter (DAT) experiments. For DAT western blot analysis, samples were not boiled before loading into the gels. Instead samples were heated at 52°C for 30 min with shaking then loaded into the gels.

### Statistical analysis

For sample size determination, we used the Sample Size Calculator at ClinCalc.com: http://clincalc.com/Stats/SampleSize.aspx. Continuous endpoint, two independent sample study design was used with the following study parameters; probability of a type-I error = 0.05, probability of a type-II error = 0.1, power = 0.9. Statistical analysis was performed using Prism software (GraphPad Software Inc.) with the alpha power level of 0.05. Unpaired *t*-test was used to perform two group comparisons. Analysis of variance (ANOVA) was used to perform multiple comparisons followed by Bonferroni’s multiple comparisons test. For ligand binding assays, curve fitting was performed using non-linear regression analysis; specific binding with Hill slope. Unless otherwise indicated, data were graphed as means +/- SEM.

## Acknowledgements

We thank L. Hester, R. Barrow, A. Snowman, S. McTeer and L. Albacarys from the SHS laboratory for their assistance. We thank T. Boronina and the Mass Spectrometry Core at Johns Hopkins School of Medicine for helping with the mass spectrometry experiments. We are also grateful for fruitful discussions with L. Mario Amzel, Ph.D., Robert N. Cole, M.S., Ph.D., members of the S.H.S. laboratory, and Deiaa Harraz.

## Author Contributions

MMH, PC, IGK, APM, ERS, AMS, MS, YJS, and YH performed experiments. MMH and SHS designed experiments, analyzed data, and wrote the manuscript.

## Author Information

Conflict of interest: The authors declare none.

Correspondence and requests for materials should be addressed to.

---

**Supplementary Figure 1.**
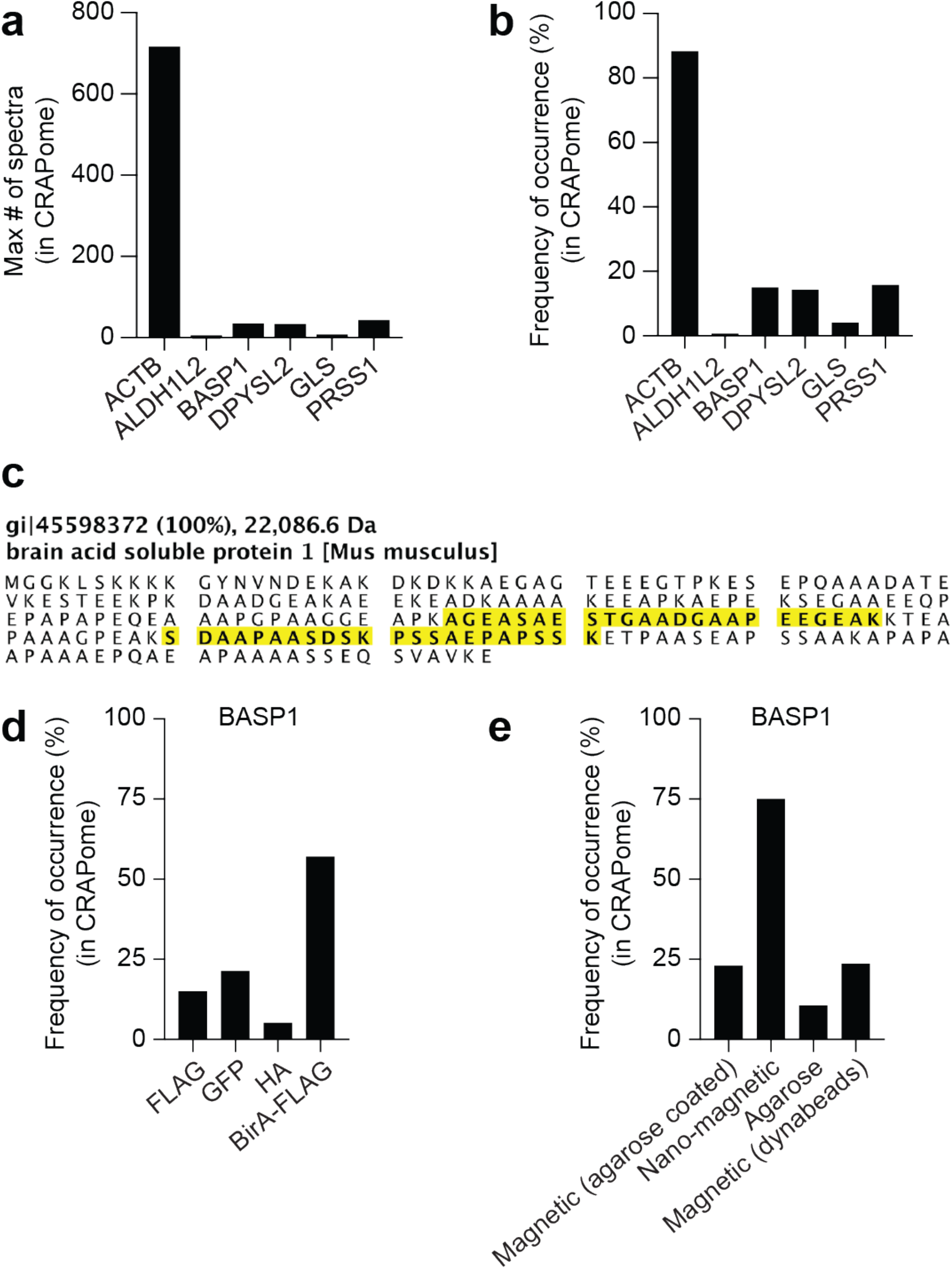
Identification of cocaine binding proteins. **a & b**, Contaminant Repository for Affinity Purification (CRAPome) search results for the six mouse proteins we identified by mass spectrometry to co-immunoprecipitate with cocaine. Maximum number of spectra for each protein (**a**) and the frequency of occurrence in all CRAPome experiments (**b**) are shown. **c**, Mass spectrometry identified BASP1 in anti-cocaine immunoprecipitates from primary cortical neurons. The identified peptides mapped to BASP1 amino acid sequence are shown. **d & e**, Distribution of identifying BASP1 across different epitope tag (**d**) and affinity support (**e**) purifications in CRAPome. BASP1 is not frequently detected, and often with a low number of spectral counts. Also, BASP1 detection is frequently associated with FLAG tag and magnetic bead pull down assays but much less so with HA tag and agarose beads. Taken together the data from the CRAPome search suggest that BASP1 is unlikely to be a common contaminant in affinity pull down/mass spectrometry studies unless associated with FLAG tag or magnetic bead pull downs.

**Supplementary Figure 2.**
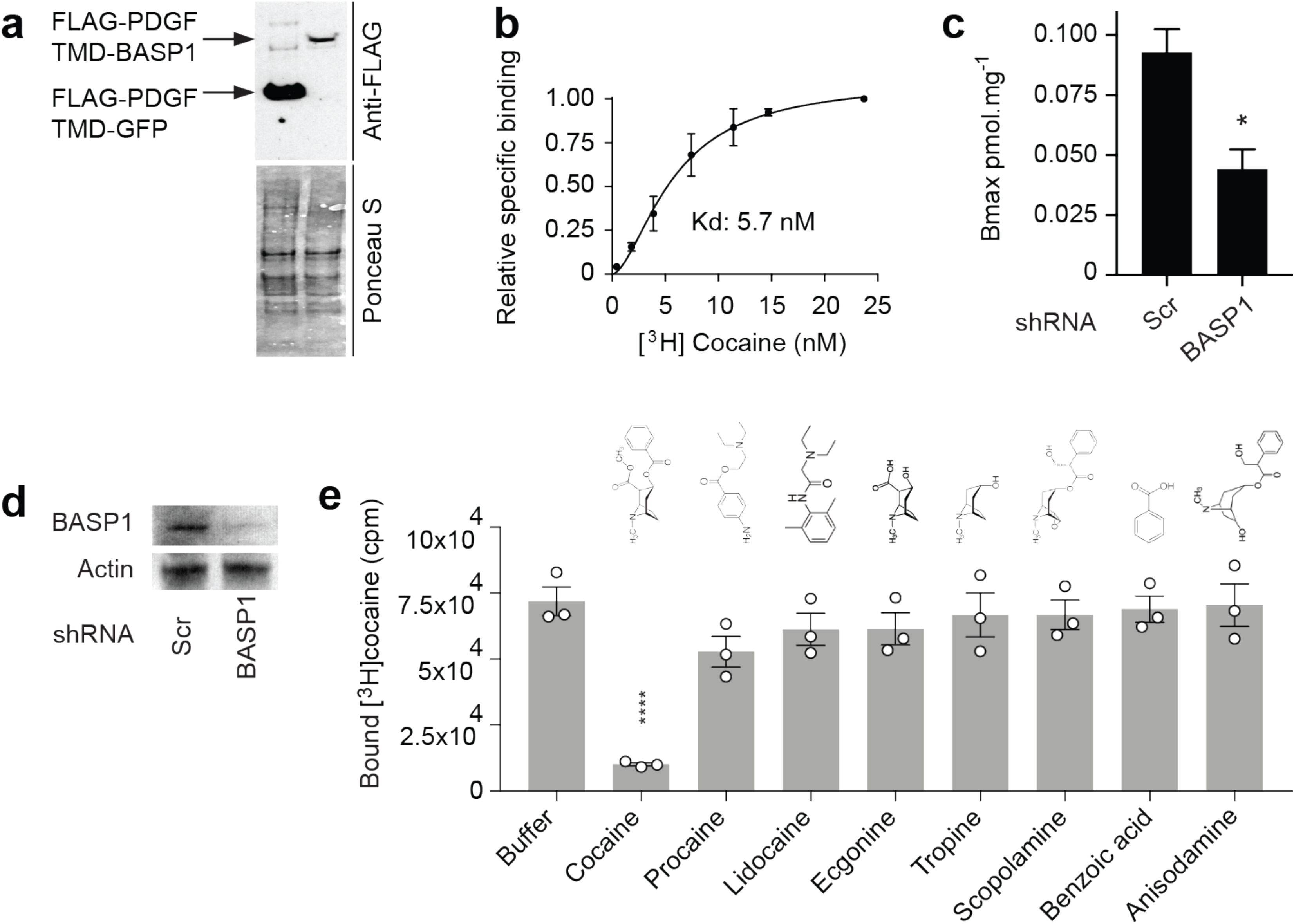
BASP1 binds [^3^H]cocaine. **a**, Western blot analysis of membrane fractions isolated from HEK 293 cells’ membrane fractions confirming the overexpression of FLAG tagged PDGF transmembrane domain (FLAG-PDGF-TMD) fused to the N-termini of GFP or BASP1. **b**, Saturation binding curve for [^3^H]cocaine and BASP1 with myc-FLAG tags on its C-terminus overexpressed in HEK-293 cells. Membrane fraction is used for the ligand binding assays. Kd = 5.8 nM, *n* = 3, R square = 0.97, specific binding with Hill slope. **c**, A histogram for the Bmax (maximum number of binding sites) for [^3^H]cocaine in synaptosome fractions isolated from rat striata following injection with AAV2 encoding Scr or BASP1 shRNA. BASP1 depletion in rat striatum diminishes the Bmax of for [^3^H]cocaine binding. *n* = 2, * (*P* = 0.0314), one-tailed *t* test. Error bars = +/-SEM. **d**, Western blot analysis of rat striatal synaptosome fractions confirming the knockdown of BASP1 protein. **e**, Specific binding of [^3^H]cocaine to membrane fraction of HEK 293 cells overexpressing BASP1 following elution with indicated structural analogues/derivatives of cocaine. *n* = 3, one-way ANOVA, *P* < 0.0001, Bonferroni’s multiple comparisons test, **** (*P* <0.0001). Error bars = +/-SEM.

**Supplementary Figure 3.**
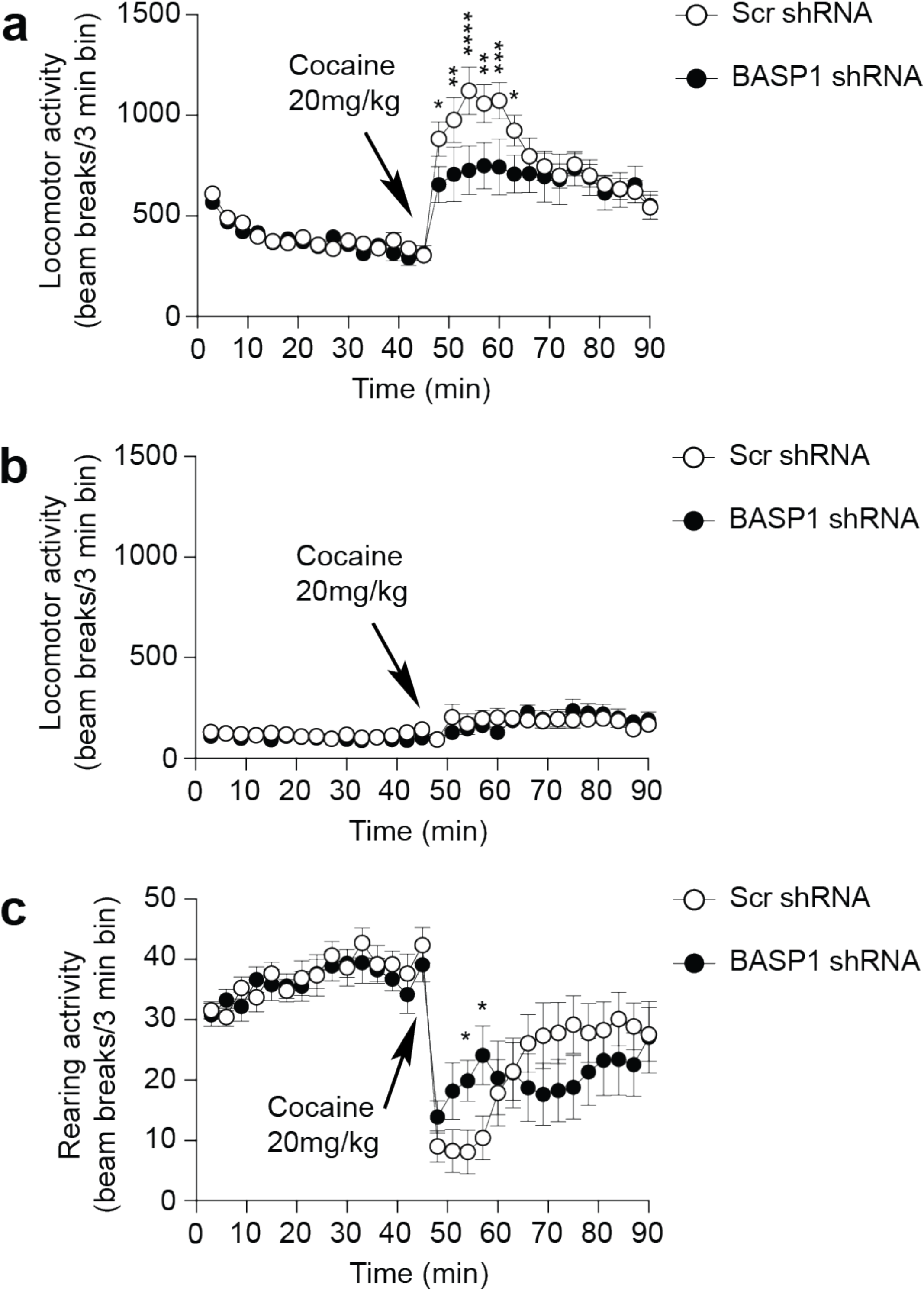
BASP1 regulation of the locomotor stimulant effect of cocaine. **a**, Peripheral locomotor activity of mice in the open field test treated with cocaine preinjected with AAV2 scrambled (Scr) or BASP1 shRNA. *n* = 11 mice/group, two-way ANOVA, * (*P* = 0.0197 and 0.0268 respectively), ** (*P* = 0.0057 and 0.0017 respectively), *** (*P* = 0.0008), **** (*P* < 0.0001). Error bars = +/- SEM. **b**, Central locomotor activity of mice in the open field test treated with cocaine preinjected with AAV2 scrambled (Scr) or BASP1 shRNA. *n* = 11 mice/group. Error bars = +/-SEM. **c**, Rearing activity of mice in the open field test treated with cocaine preinjected with AAV2 scrambled (Scr) or BASP1 shRNA. *n* = 11 mice/group. two-way ANOVA, * (*P* = 0.03 and 0.012 respectively). Error bars = +/-SEM.

**Supplementary Figure 4.**
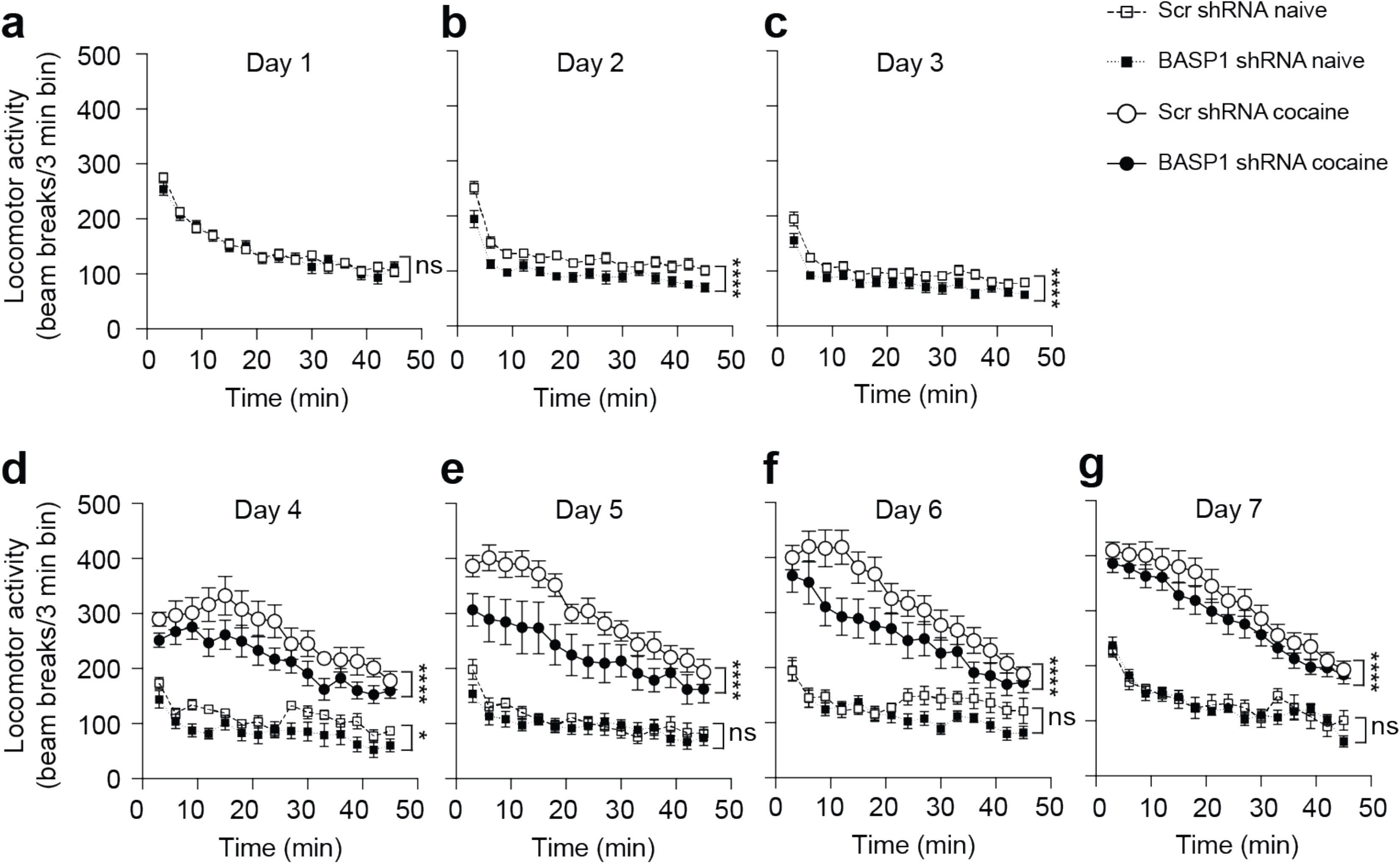
BASP1 regulates locomotor sensitization to cocaine. **a-g**, Daily total locomotor activity of mice in the open field test treated with saline or cocaine (15 mg/kg), preinjected with AAV2 scrambled (Scr) or BASP1 shRNA. Scr shRNA naive group: *n* = 7, BASP1 shRNA naive group: *n* = 9, Scr shRNA cocaine group: *n* = 16, BASP1 shRNA cocaine group: *n* = 13. Two-way ANOVA, Bonferroni’s multiple comparisons test, * (*P* = 0.038), **** (*P* < 0.0001). Error bars = +/-SEM.

